# Phase variable colony morphotypes of *Clostridioides difficile* elicit distinct host responses during acute infection

**DOI:** 10.64898/2026.01.09.698580

**Authors:** Jilarie A. Santos-Santiago, Nicole C. Gadda, Rita Tamayo

## Abstract

Many *Clostridioides difficile* strains can form two colony morphotypes: rough and smooth. The rough and smooth morphotypes differ in multiple phenotypes, including cell length and chaining, motility, biofilm production, and virulence in the hamster model of *C. difficile* infection (CDI). Colony morphology undergoes phase variation and is determined by the ON/OFF expression of *cmrRST*, which encodes a signal transduction system. Here, we test the hypothesis that differences in colony morphology and the associated phenotypes influence pathogenesis and the host response to infection. We first compared the rough and smooth colony variants of wild-type *C. difficile* in a mouse model of CDI. However, CmrRST phase varied during infection such that the *C. difficile* populations became indistinguishable in feces and tissues, and no differences in disease were observed in mice inoculated with these variants. We next circumvented phase variation using mutants that form only rough or only smooth colonies. Co-infection of mice with these phenotypically locked strains revealed that the smooth colony mutant has greater fitness than the rough mutant, which is outcompeted during late infection. In addition, NanoString analyses showed a higher number of differentially expressed pro-inflammatory genes and overall higher expression levels in mice infected with the rough colony mutant, independent of bacterial burden and toxin levels. Our results indicate that in a mouse model of CDI, cells from rough colonies are more immunostimulatory during early murine infection, potentially leading to reduced relative fitness compared to cells from smooth colonies.

**IMPORTANCE:** *Clostridioides difficile* is a major cause of antibiotic-associated diarrhea and one of the most common hospital-acquired infections. These infections are often recurrent and recalcitrant to antibiotic treatment. Bacteria use diverse mechanisms to adapt to stressful host environments, including the development of different subpopulations to help ensure survival of the whole population. *C. difficile* produces two different colony types, rough and smooth, that have multiple distinguishing traits. We interrogated the effect of these different forms of *C. difficile* on pathogenesis and the host immune response. We show that the rough colony phenotype results in a more robust immune response and has reduced fitness during infection compared to the smooth colony phenotype. This work sheds light on the impact of these phenotypic subpopulations on disease, leading to a deeper understanding of *C. difficile*-host interactions.

## INTRODUCTION

*Clostridioides difficile* is a Gram-positive, spore-forming bacterium and one of the leading causes of nosocomial infections worldwide (1,2). *C. difficile* ingestion can result in asymptomatic intestinal carriage, diarrhea, and more severe conditions such as pseudomembranous colitis (3). *C. difficile* infection (CDI) is associated with antibiotic exposure that disrupts the intestinal microbiota and generates an environment conducive to *C. difficile* growth and production of the glucosylating toxins TcdA and TcdB, which are essential drivers of disease development (4,5). The host immune response to toxin-induced damage and other *C. difficile* virulence factors contributes to the severity of CDI. The immune response involves increased production of pro-inflammatory cytokines and chemokines (i.e., IL-1β, IL-6, CXCL1, CXCL2, and TNF-α) that then aid in immune cell recruitment for pathogen clearance (6–10). Cytokine and chemokine profiles have been investigated as markers for CDI severity independent of bacterial burden (11,12). Recent work demonstrated a significant increase in the serum levels of IL-1β, IL-6, and TNF-α in CDI patients compared to healthy individuals (12). In addition, neutralization of CXCL1 and CXCL2 during early CDI in mice improved host survival and decreased the neutrophil count in peripheral blood (8).

During infection, *C. difficile* colonizes the large intestine, which normally harbors the diverse bacterial microbiota crucial for suppressing *C. difficile* spore germination and vegetative cell growth (16). The large intestine provides distinct nutritional niches that can be altered during enteric infections (13–15). Bacteria can employ a variety of mechanisms to adapt to these niches, including the development of phenotypically distinct subpopulations that can differ in fitness depending on the environment (16–18). This phenomenon has been investigated in other pathogenic bacteria, including *Streptococcus pneumoniae*, which produces transparent and opaque colony variants associated with nasopharyngeal colonization and sepsis, respectively (19). In another case, *Escherichia coli* controls the expression of type I fimbrial genes through an invertible promoter, where one orientation leads to fimbrial gene transcription and enhances bacterial colonization in the kidney and bladder (20). The *E. coli* population with the invertible promoter in the opposite orientation is non-fimbriated and is the main variant found in urine isolates (20,21).

Extensive work has demonstrated that *C. difficile* develops phenotypically heterogeneous populations (22–29). Many *C. difficile* strains, including the epidemic-associated strain R20291, form two distinct colony morphologies: rough colonies surrounded by tendril-like structures or smooth colonies with rounded edges (26,30). *C. difficile* can reversibly switch between rough and smooth colony phenotypes through phase variation of the *cmrRST* operon, which encodes two response regulators, CmrR and CmrT, and a histidine kinase, CmrS (26,31,32). We previously demonstrated that CmrT is required for rough colony formation *in vitro* (26,33), while CmrR positively autoregulates transcription of *cmrRST* (31). Expression of *cmrRST* is controlled in a binary manner by the orientation of an invertible DNA element, the *cmr* switch, upstream of the operon (25,26,31). The *cmr* switch contains a promoter such that one orientation of the switch leads to *cmrRST* transcription and rough colony formation (*cmr-*ON); in the opposite orientation, the promoter is directed away from *cmrRST*, so this transcription is lost and smooth colonies develop (*cmr*-OFF) (31). Further characterization revealed additional differences *in vitro* between the colony morphotypes. The rough colony morphology displays increased cell length and chaining, greater surface motility, reduced swimming motility, and reduced biofilm production relative to the smooth morphotype (26,31).

In a hamster model of acute CDI, a rough colony isolate showed greater virulence when compared to a smooth colony isolate of the same genetic background despite comparable toxin accumulation (26). Notably, phase variation of CmrRST occurred during infection— *C. difficile* populations in the cecal lumen of moribund hamsters showed a bias toward the *cmr*-OFF state regardless of whether the inoculum was *cmr*-ON (rough) or *cmr*-OFF (smooth) (26). These results suggest a toxin-independent role for CmrRST phase variation in the severity of CDI. However, hamsters are highly sensitive to TcdA and TcdB and develop a rapid and fatal disease when infected with *C. difficile*, which is not characteristic of the majority of human CDI (34). In this work, we used a mouse model of CDI, which better recapitulates typical human infection, to evaluate the host immune responses to the rough and smooth colony variants. Our results indicate that *C. difficile* from rough colonies elicits a more robust immune response and shows reduced intestinal colonization compared to cells from smooth colonies.

## RESULTS

### Comparison of wild-type rough and smooth colony variants in a murine model of CDI

To examine the impact of colony morphology and the associated phenotypes on disease, we tested previously isolated and characterized rough and smooth variants of wild-type (WT) *C. difficile* R20291 in a mouse model of CDI (26,35,36). C57BL/6 mice inoculated with spores of each variant were monitored daily for disease symptoms by measuring weight loss, and fecal samples were collected daily for assessment of bacterial burden. Mock-inoculated mice were included as controls. All inoculated mice began to lose weight by day 1 post-inoculation (p.i.). Both groups showed the highest weight loss on day 2 p.i., and the mice recovered weight thereafter (**Figure 1A**). No significant differences in weight loss or *C. difficile* burden in feces were found between rough- and smooth-inoculated mice at any time point (**Figure 1B, Fig. S1**).

**Figure 1.**
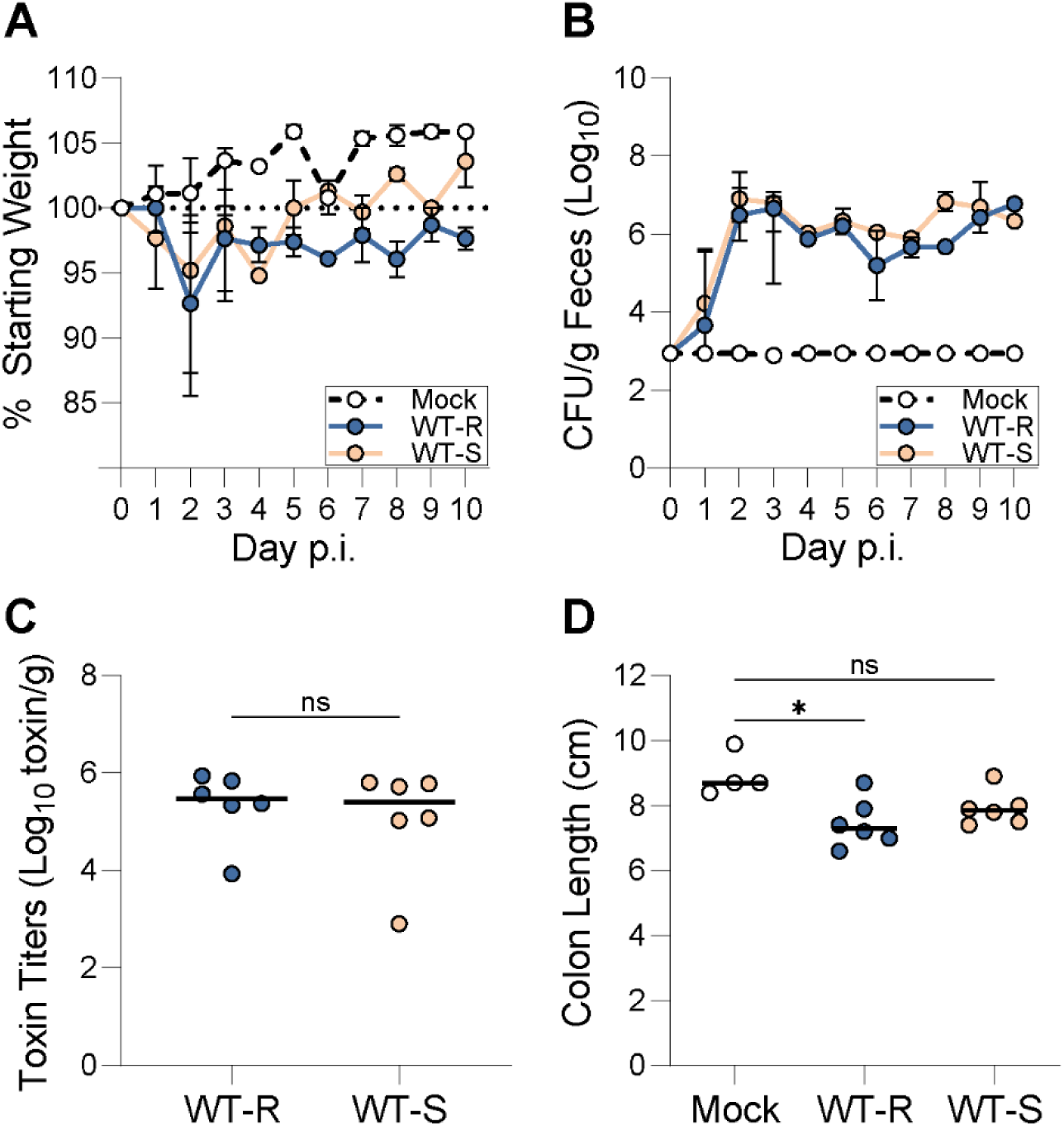
Analysis of rough and smooth colony variants of *C. difficile* in a murine model of CDI. Antibiotic-treated mice were inoculated with spores from *C. difficile* R20291 wild-type isolates displaying a rough (WT-R) or smooth (WT-S) colony morphology. Two independent experiments were conducted, and the data were pooled for analysis. **(A)** *C. difficile*-induced weight loss over time. Animal weights were measured daily post-inoculation (p.i.) and expressed as a percentage of each mouse’s starting weight at the time of inoculation (day 0). **(B)** *C. difficile* burden in feces over time. *C. difficile* colony-forming units (CFU) were enumerated from fecal samples collected daily. The dotted line represents the limit of detection. No *C. difficile* was detected in the mock-inoculated mice. **(A-B)** Circles indicate medians, and error bars indicate the range. **(C)** Toxin levels in cecal contents on day 2 p.i.. Toxin titers were calculated as the reciprocal of the highest dilution that caused ≥ 80% rounding of Vero cells. No cell rounding occurred when treated with cecal contents from mock-inoculated animals. **(D)** Colon length on day 2 p.i. measured in centimeters (cm). **(C-D)** Circles represent values from individual animals, lines indicate medians. * p ≤ 0.05 and ns = not significant; **(A-B, D)** Kruskal-Wallis test with Dunn’s post-test or **(C)** Mann-Whitney test.

We also evaluated the rough and smooth colony variants for toxin production and colon length, which serves as a proxy for intestinal inflammation (37). In murine CDI, colon shrinkage has been linked to toxin production and activity (38). Subsets of mice from each group were euthanized at the peak of infection (day 2 p.i.), the large intestines were harvested and measured, and cecal contents were collected and assayed for toxin levels. Although no significant differences in toxin were detected between inoculated groups (**Figure 1C**), we observed significant colonic shrinkage in mice inoculated with the rough, but not smooth, colony variant when compared to mock (**Figure 1D**). These results suggest that the rough colony variant caused somewhat greater inflammation independent of toxin levels and bacterial burden.

### Wild-type rough and smooth colony variants undergo phase variation early during infection

Given the modest differences observed and prior results indicating that CmrRST phase variation occurs during infection of hamsters (26), we examined phase variation dynamics in the mouse model. As a proxy for the rough and smooth colony phenotypes present in the gut, we used quantitative PCR (qPCR) to determine the proportion of *C. difficile* with the *cmr* switch in each orientation in fecal samples compared to the spore inoculum. As expected, the spore inoculums (sp) from the rough colony variant had the *cmr* switch predominantly in the ON orientation (94.7% *cmr*-ON) (**Figure 2A**), while smooth colony-derived spores were mostly *cmr-*OFF (4.0*% cmr*-ON) (**Figure 2D**). In the feces of mice inoculated with the rough colony variant, we observed increased heterogeneity in *cmr* switch orientation in the *C. difficile* populations, with a decrease in the proportion of *cmr*-ON cells on day 1 p.i. This heterogeneity persisted on day 2 p.i., after which the populations shifted to a significant *cmr-*OFF bias by day 3 p.i. (**Figure 2A**). In contrast, in the smooth variant-inoculated mice, *cmr* switch inversion to the *cmr*- ON orientation occurred in some mice by day 1 p.i.; however, the effect was transient, and there were no significant differences in the *cmr*-ON/OFF make-up of these fecal populations compared to the inoculum (**Figure 2D**). Overall, for both rough and smooth variant inoculums, the populations in feces were heterogeneous on days 1 p.i. and skewed *cmr*-OFF by day 3 p.i. (**Figure 2A, D**). These results suggest that regardless of the starting orientation of the *cmr* switch, the populations in feces rapidly converge to a *cmr*-OFF state.

**Figure 2.**
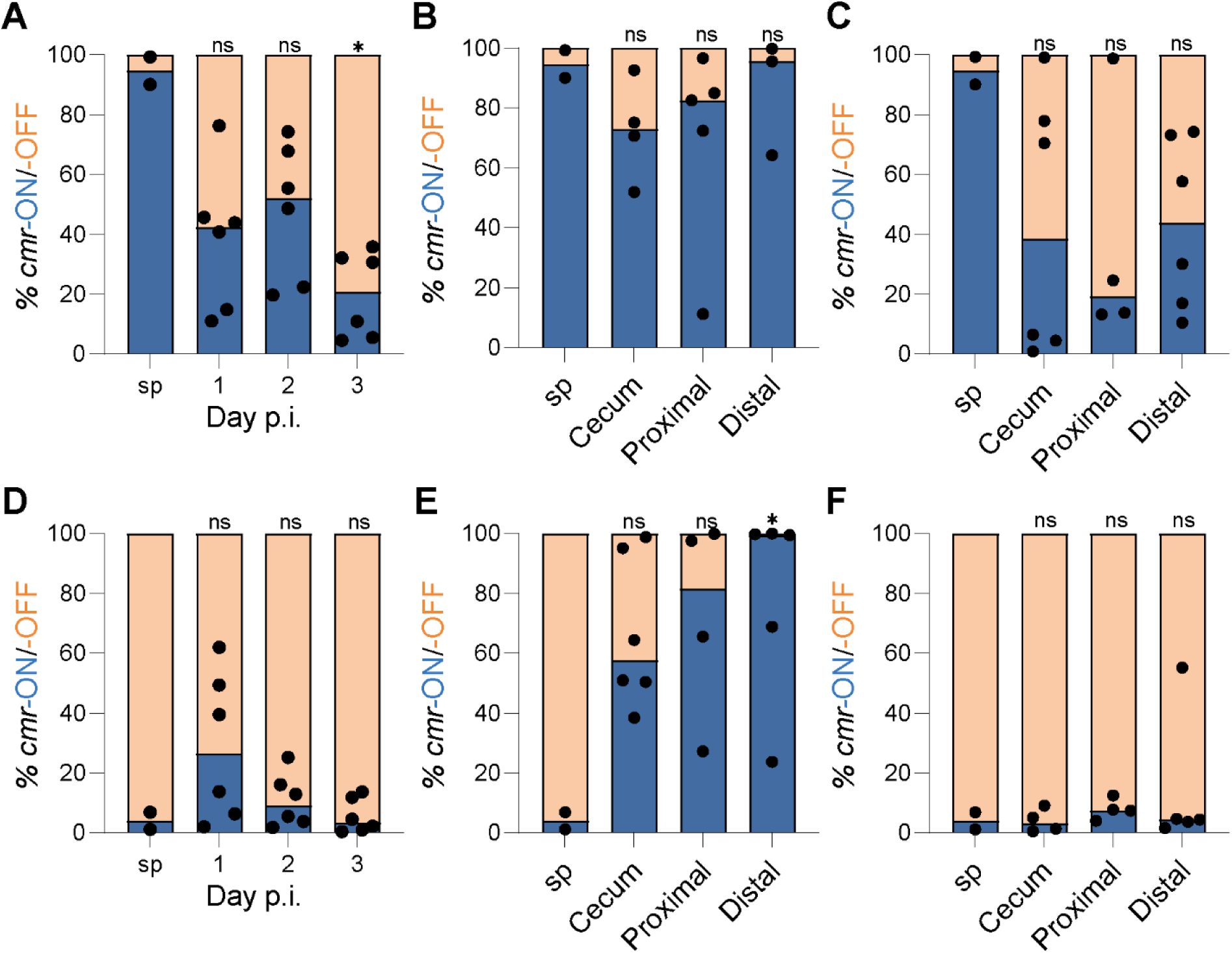
Orientation of the *cmr* switch in tissue-associated *C. difficile* during early stages of infection. Mice were inoculated with spores from wild-type *C. difficile* isolates displaying a rough (WT-R) **(A-C)** or smooth (WT-S) **(D-E)** colony morphology. DNA was purified for the determination of *cmr* switch orientation by qPCR from feces on days 1, 2, and 3 p.i. **(A, D)**, or intestinal tissue divided into cecum, proximal colon, and distal colon on day 1 p.i. **(B, E)** or day 2 p.i. **(C, F)**. DNA from the spore inoculums (sp) was analyzed for comparison to the starting populations. Inoculums from panels A-C and D-F were collected from the same experiment, respectively. Data are expressed as the percentage of the population with the *cmr* switch in the ON (*cmr*-ON, blue) or OFF (*cmr*-OFF, orange) orientation. Circles represent values from individual animals, bars indicate medians. *** p ≤ 0.001, ** p ≤ 0.01, * p ≤ 0.05, and ns = not significant; Kruskal-Wallis test with Dunn’s post-test in comparison to inoculum.

Because the *cmr*-ON/OFF makeup of *C. difficile* populations in feces may not reflect the populations retained in the intestinal tract, we also examined the proportions of *cmr*-ON/OFF cells associated with tissues. Groups of mice inoculated with the rough or smooth colony variant were euthanized on days 1 and 2 p.i., and large intestines were harvested and divided into cecum, proximal, and distal colon segments for qPCR analysis of *cmr* switch orientation. We found that the *C. difficile* populations in all intestinal segments from mice inoculated with the rough colony variant maintained a *cmr*-ON bias on day 1 p.i. (**Figure 2B**). By day 2 p.i., a qualitative shift to *cmr*-OFF was observed (**Figure 2C**). In the mice inoculated with the smooth colony variant, the *C. difficile* populations in all tissue segments showed greater heterogeneity on day 1 p.i., with a significant increase in the proportion of *cmr*-ON cells in the distal colon compared to the spore inoculums (**Figure 2E**). Interestingly, by day 2 p.i., the populations were almost uniformly *cmr*-OFF in all tissue segments (**Figure 2F**).

The differences in proportions of *cmr-*ON/OFF cells present in feces and tissues were independent of bacterial burden, as the colony-forming units (CFU) were comparable in rough- and smooth-inoculated mice (**Fig S2**). Taken together, these results indicate that phase variation of CmrRST is dynamic during early infection, with selective enrichment of *cmr-*ON or *cmr*-OFF cells based on time post-inoculation and intestinal niche. Specifically, regardless of whether the inoculum is *cmr*-ON or *cmr*-OFF, *cmr-*ON cells tend to dominate in tissue-associated populations of *C. difficile* on day 1 p.i., in contrast to fecal populations, which showed a shift to the *cmr-*OFF state within the same time frame. By day 2 p.i., tissue and fecal *C. difficile* populations both shifted to a *cmr-*OFF state.

### Effect of CmrRST on the overall colonic inflammation during early infection

In infections with WT rough and smooth colony variants, the *cmr* switch orientation quickly became indistinguishable, complicating the investigation of the host response to the different colony morphotypes. To circumvent this hurdle, we used mutants that are phenotypically locked : a *cmr*Δ3-ON mutant with the *cmr* switch locked in the ON orientation that produces only rough colonies *in vitro*, and a Δ*cmrR*Δ*cmrT* mutant lacking the CmrR and CmrT regulators that produces only smooth colonies (**Figure 3A**) (31). We opted to use the deletion mutant instead of a phase-locked OFF mutant, because *cmrRST* transcription can occur via other, independent mechanisms (31,66). Mice were inoculated with 10^5^ spores of the mutants or of WT *C. difficile*, where our freezer stock of the WT is predominantly *cmr*-OFF, but *cmr* switch inversion to *cmr*-ON can occur (26,31). The orientation of the *cmr* switch in the spore inoculums was confirmed before inoculation (**Fig. S3**). The mice were assessed for weight loss, bacterial burden, colon length, and toxin accumulation.

**Figure 3.**
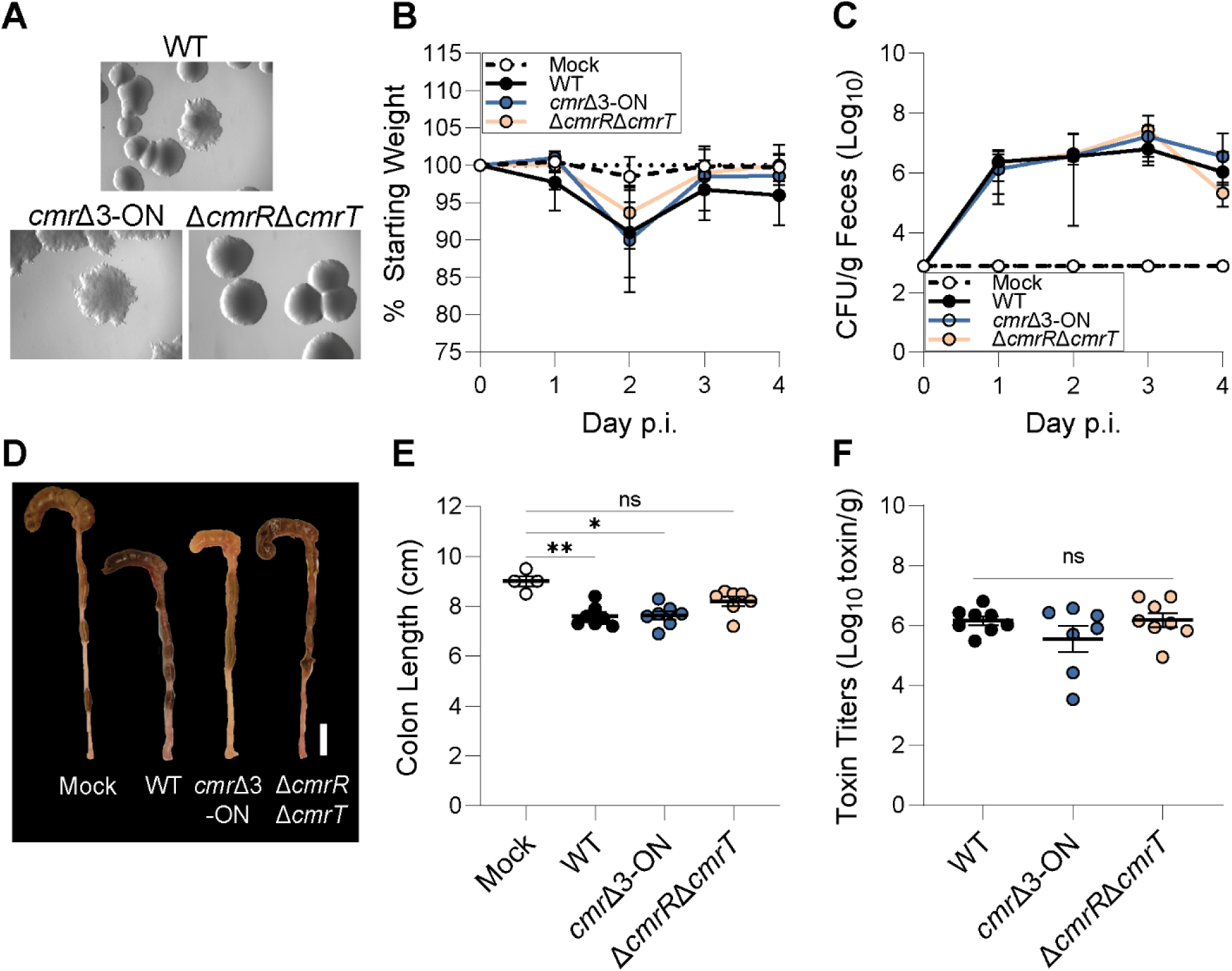
Assessment of phenotypically locked rough and smooth mutants in murine CDI. Mice were inoculated with spores of WT, *cmr*Δ3-ON, or Δ*cmrR*Δ*cmrT*. Two independent experiments were conducted, and the data were pooled for analysis. **(A)** Representative images of colony morphology of WT, *cmr*Δ3-ON, and Δ*cmrR*Δ*cmrT* strains after 48 hours of growth on BHIS-agar. Images were taken at 2X magnification. **(B)** *C. difficile*-induced weight loss over time. Animal weights were measured daily p.i. and expressed as a percentage of each mouse’s starting weight at the time of inoculation (day 0). **(C)** *C. difficile* burden in feces over time. *C. difficile* colony-forming units (CFU) were enumerated from fecal samples collected daily. The dotted line represents the limit of detection. No *C. difficile* was detected in the mock-inoculated mice. **(D)** Representative image of the large intestines removed on day 2 p.i. Scale bar = 1 cm. **(E)** Quantification of colon length. Colons were harvested on day 2 p.i., and lengths were measured in cm. **(F)** Toxin levels in the cecal contents. Toxin titers were calculated as the reciprocal of the highest dilution to cause ≥ 80% rounding of Vero cells. No cell rounding occurred when treated with cecal contents from mock-inoculated animals. **(B-C)** Circles indicate medians, and error bars indicate the range. **(E-F)** Circles represent values from individual animals, and lines indicate medians. ** p ≤ 0.01, * p ≤ 0.05, and ns=not significant; Kruskal-Wallis with Dunn’s post-test.

On day 1 p.i., mice infected with WT showed small but statistically significant weight loss when compared to mock and the mutant-inoculated groups (**Figure 3B**, **Fig. S4**). By day 2 p.i., all inoculated groups showed statistically significant weight loss when compared to mock, but there were no differences among the infected groups (**Figure 3**, **Fig. S4**). The weight loss of the infected groups was comparable to that of the mock group by day 3 p.i., suggesting recovery. No significant differences in *C. difficile* burden were detected at any time point (**Figure 3C**, **Fig. S4**). The colons from mice inoculated with WT and *cmr*Δ3-ON were significantly shorter than colons from the mock controls (**Figure 3D – E**). Colon lengths from the Δ*cmrR*Δ*cmrT* inoculated mice were not different from the mock controls, indicating different inflammation levels in the colon between the two mutants. Differences in colon length were not due to toxin levels, as no significant differences in cecal toxin accumulation were detected between any of the inoculated groups (**Figure 3F**). These results suggest a role for CmrRST in overall colonic inflammation during early infection.

### The phase-locked smooth colony mutant outcompetes the locked rough mutant during infection

The differences in pathogenesis in mice infected with phenotypically locked rough and smooth colony mutants were subtle, but the observed phase variation occurring during infection (Figure 2) suggests that cells from rough and smooth colony variants differ in fitness. Specifically, the shift of *C. difficile* to a predominantly *cmr*-OFF population suggests the smooth colony variant is better adapted to the intestinal environment. To directly test this idea, we conducted competition experiments with the phenotypically locked mutants in mice. To allow differentiation of the two strains, we integrated a spectinomycin resistance cassette (*aad9*) at a neutral site in the *cmr*Δ3-ON chromosome, creating the marked strain *cmr*Δ3-ON^R^. After co-inoculation with a 1:1 ratio of spores from *cmr*Δ3-ON^R^ and Δ*cmrR*Δ*cmrT*, fecal samples were collected daily for enumeration of CFU on agar medium with or without spectinomycin. We then calculated the competitive index between the mutants in addition to determining the bacterial load for each strain. *In vitro* co-culture indicates comparable fitness between *cmr*Δ3-ON^R^ and Δ*cmrR*Δ*cmrT* in rich medium (**Fig. S5A**).

For the first four days p.i., the median competition index remained ∼ 1, indicating no differences in fitness during this time frame (**Figure 4A**). From day 5 p.i. and thereafter, the competitive indices significantly decreased, with Δ*cmrR*Δ*cmrT* outcompeting *cmr*Δ3-ON^R^. By day 6 p.i., the *cmr*Δ3-ON^R^ mutant was mostly cleared while the Δ*cmrR*Δ*cmrT* burden remained consistent and reached significantly higher density than *cmr*Δ3-ON^R^ (**Figure 4B**). The clearance of *cmr*Δ3-ON^R^ was not due to the insertion of *aad9*—the selectable marker was introduced at the same chromosomal site in the WT background, and co-infection of mice along with the unmarked WT parent strain indicated no difference in fitness due to the marker throughout the same time-frame (**Fig. S5B**). These results are consistent with the observed shift to a *cmr*-OFF population in the feces of WT-infected mice (Figure 2) and suggest that the *cmr*-ON state becomes disadvantageous at later time points during infection.

**Figure 4.**
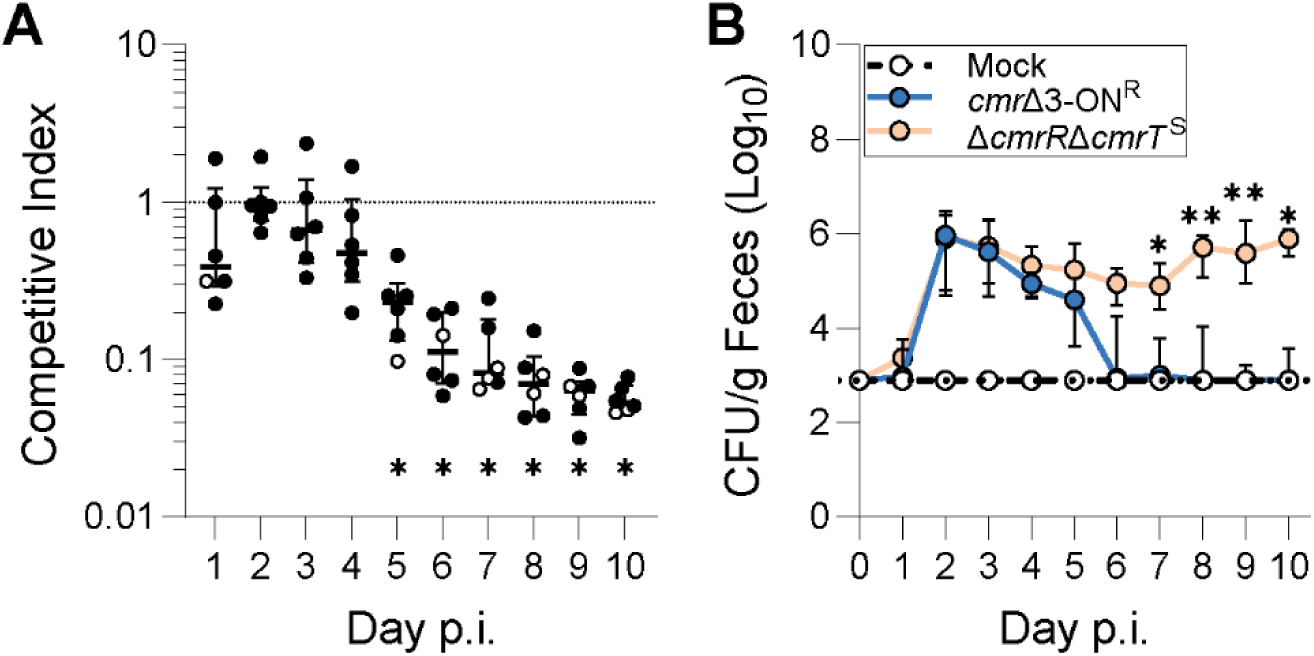
Phenotypically locked smooth outcompetes locked rough mutant during later stages of infection. Mice were inoculated with a mixed inoculum of a 1:1 ratio of *cmr*Δ3-ON^R^ and Δ*cmrR*Δ*cmrT* spores. Feces were collected daily and plated on TCCFA (total spores) and TCCFA with spectinomycin (*cmr*Δ3-ON^R^ spores only) to enumerate CFU of each strain and calculate the competitive index (CI). Two independent experiments were conducted, and the data were pooled for analysis. **(A)** Relative fitness expressed as a CI. Circles represent CI values from the feces of individual animals. Open circles are approximate CI values where Δ*cmrR*Δ*cmrT* was below the limit of detection (LoD). **(B)** Burden of the individual mutants in feces over time. The dotted line represents the LoD. No *C. difficile* was detected in the mock-inoculated mice. Circles indicate medians, and error bars indicate the range. ** p ≤ 0.01, and * p ≤ 0.05; (A) Wilcoxon rank sum test comparing values to a hypothetical CI of 1 indicating equal fitness, and (B) Kruskal-Wallis test with Dunn’s post-test.

### Phenotypically locked smooth *C. difficile* elicits lower expression of *Il1β*, *Cxcl1*, *Lcn2*, and *Tnfα* in mice

The competitive disadvantage of the phenotypically locked rough *cmr*Δ3-ON^R^ mutant compared to the smooth Δ*cmrR*Δ*cmrT* mutant could be driven by the host environment. We therefore evaluated the host response to these mutants and WT *C. difficile* in the mouse model. The ceca from the infected mice and mock-inoculated controls were collected at the peak of infection (day 2 p.i.), and RNA was extracted for qRT-PCR analysis of the abundance of transcripts encoding the pro-inflammatory cytokines *Il1β*, *Cxcl1*, *Lcn2,* and *Tnfα*. The production of these cytokines has been associated with *C. difficile* infection and neutrophil recruitment to the site of infection (12,39–41).

Compared to the mock controls, mice infected with WT or *cmr*Δ3-ON showed a significant Log_2_ fold increase (3.02- and 3.34, respectively) in *Il1β* transcript, while Δ*cmrR*Δ*cmrT* resulted in a non-significant 1.26-fold change (**Figure 5A**). For *Cxcl1*, WT and *cmr*Δ3-ON stimulated Log_2_ fold increases of ∼ 4.58 and ∼ 5.27, respectively; the Δ*cmrR*Δ*cmrT*-inoculated mice showed a Log_2_ fold change of ∼ 1.41 (**Figure 5B**). Similarly, *Lcn2* transcript levels in WT- and *cmr*Δ3-ON-inoculated mice were Log_2_ 6.86- and 6.11-fold higher, respectively, while Δ*cmrR*Δ*cmrT*-inoculated mice showed a non-significant ∼ 2.95-fold change (**Figure 5C**). Lastly, *Tnfα* expression was Log_2_ 4.30- and 4.83-fold higher in mice inoculated with WT- and *cmr*Δ3-ON, respectively, whereas Δ*cmrR*Δ*cmrT*-inoculated mice showed a Log_2_ 2.58-fold increase (**Figure 5D**). These results suggest that infection with rough colony *C. difficile* (the *cmr*Δ3-ON mutant or WT containing the *cmr*-ON variant) elicits a stronger immune response than infection with the Δ*cmrR*Δ*cmrT* (or *cmr*-OFF) mutant.

**Figure 5.**
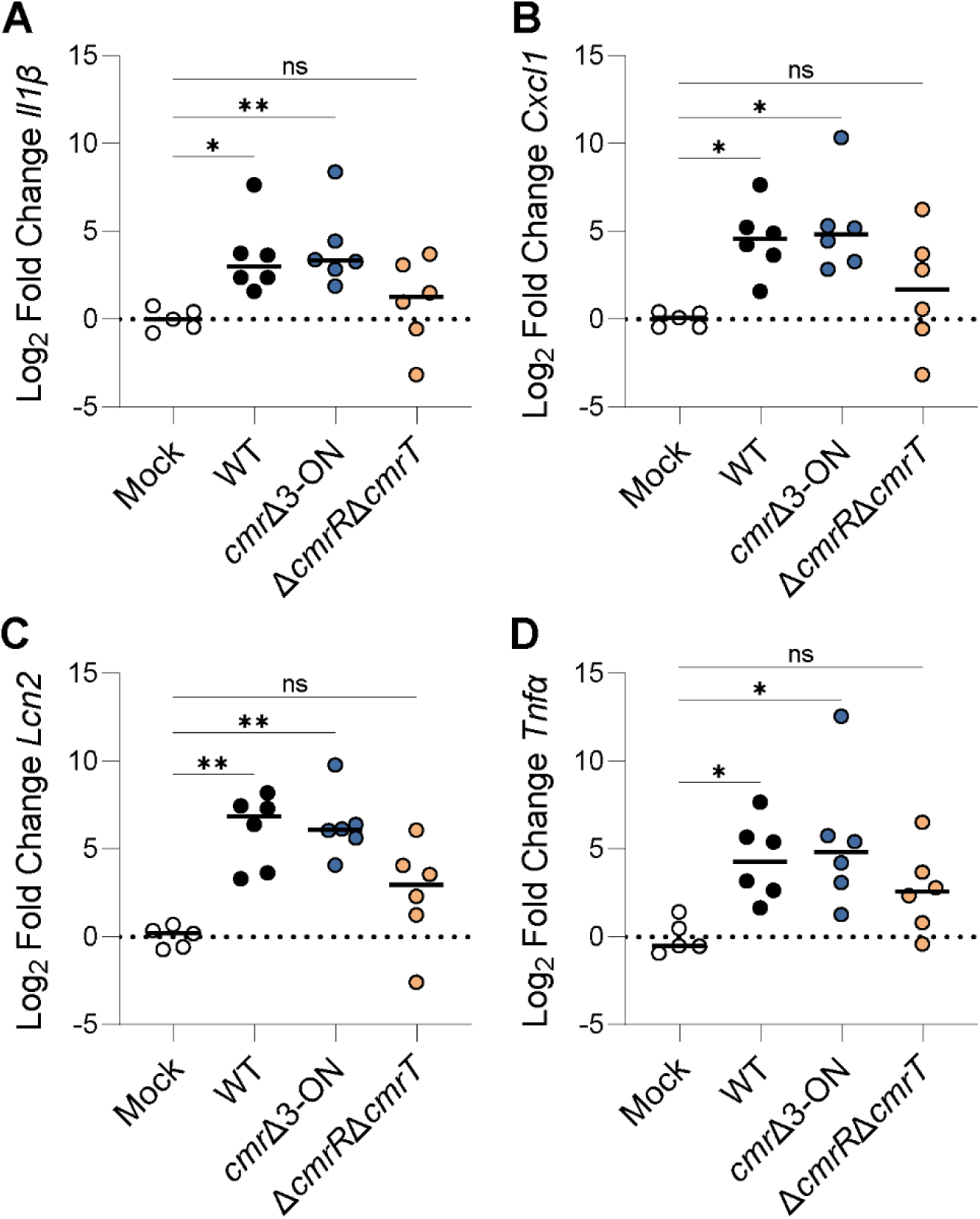
Phenotypically locked rough *C. difficile* induces greater expression of *Il1β*, *Cxcl1*, *Lcn2,* and *Tnfα* in mice. Log_2_ fold changes of **(A)** *Il1β*, **(B)** *Cxcl1*, **(C)** *Lcn2,* and **(D)** *Tnfα*. Transcription levels were determined by qRT-PCR on cDNA synthesized from RNA isolated from cecal tissue on day 2 p.i. Data were analyzed with *Tbp* as the host reference, and values were normalized to the average value of the mock samples. ** p ≤ 0.01, * p ≤ 0.05, and ns = not significant; Kruskal-Wallis test and Dunn’s post-test.

### The transcriptional response against phenotypically locked rough and smooth mutants

Based on the differences in pro-inflammatory gene expression, we conducted a broader analysis of the transcriptional changes in mice inoculated with rough and smooth colony mutants. We collected the ceca from mice inoculated with spores from WT, *cmr*Δ3-ON, or Δ*cmrR*Δ*cmrT*, as well as mock-inoculated controls, on day 2 p.i. The transcriptional profiles were evaluated using the Nanostring Mouse Host Response Panel. Infected groups were compared to the mock controls, and differential expression was determined using cut-offs of Log_2_ fold changes ≥ 1 or ≤ −1 and an adjusted p-value ≤ 0.05. This analysis showed that inoculation with WT resulted in 56 differently expressed (DE) genes (13 decreased, 43 increased) (**Figure 6A**). Mice inoculated with *cmr*Δ3-ON showed almost twice the number of DE genes (24 decreased, 76 increased; 100 total) (**Figure 6B**). Inoculation with Δ*cmrR*Δ*cmrT* yielded 47 DE genes (17 decreased, 30 increased) (**Figure 6C**). All groups shared 31 DE genes, where 22 were increased, and 9 were decreased compared to the mock controls (**Figure 6D – E**). Among these genes, the fold changes in gene expression were greater in *cmr*Δ3-ON inoculated mice than in the WT and Δ*cmrR*Δ*cmrT* groups (**Figure 6F**). The WT- and Δ*cmrR*Δ*cmrT-*inoculated mice showed 2 unique DE genes, with 1 increased and 1 decreased in each group (**Figure 6D – E**). The *cmr*Δ3-ON inoculated mice showed a higher number of unique DE genes, with 27 increased and 7 decreased. A number of these genes (*Casp3*, *Il13ra1*, *Ccl7*, *Cebpb*, *Ptgs2*, *Hsp90b1*, *Jun*, and *Nfkbia*) are involved in the IL-17 signaling pathway previously shown to affect CDI severity (42). Taken together, these results suggest that infection with the rough colony-forming *cmr*Δ3-ON mutant elicits a more robust and distinct immune response than the other *C. difficile* strains.

**Figure 6.**
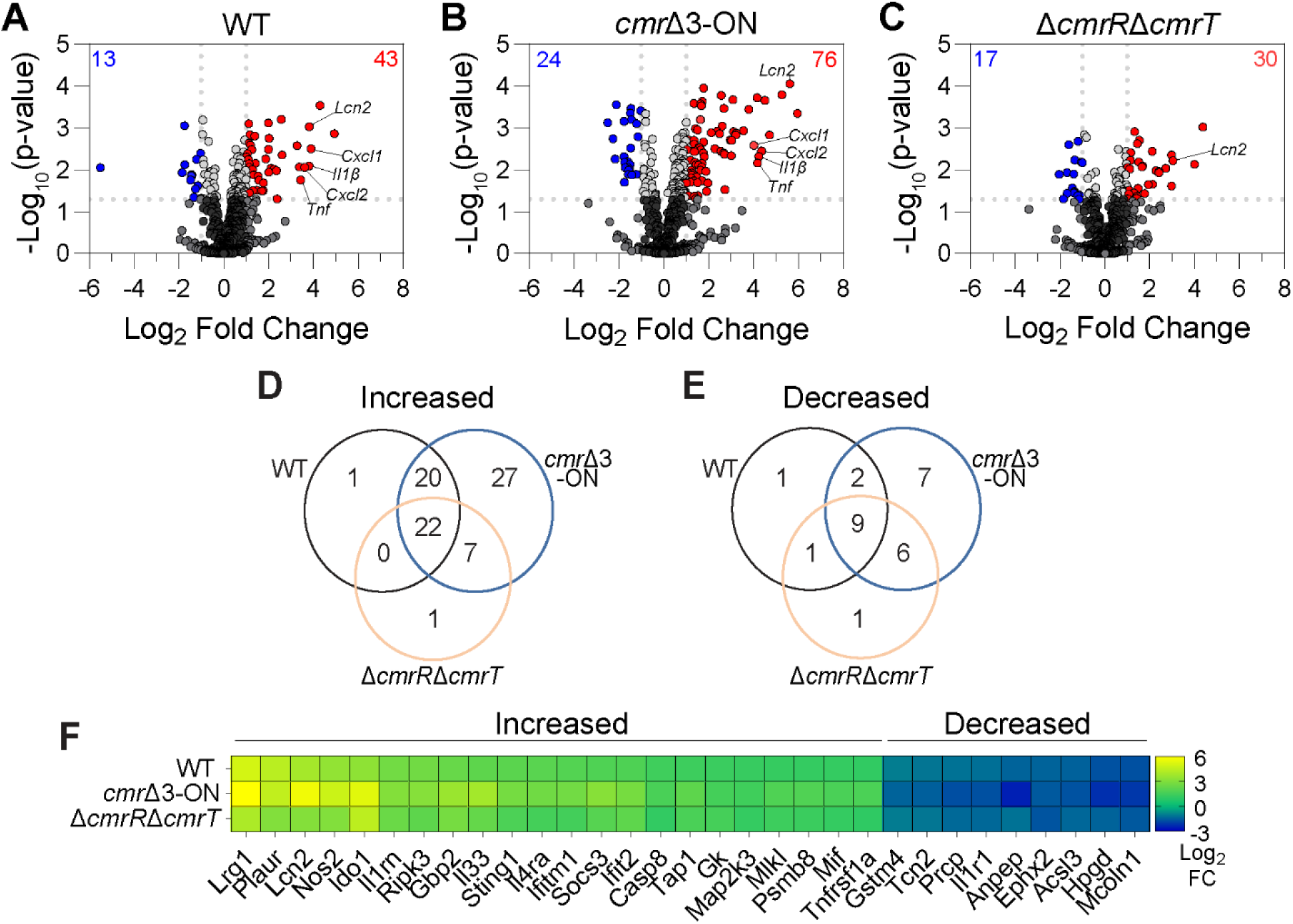
Phenotypically locked mutants alter the pro-inflammatory transcriptional response at the peak of infection. Mice were inoculated with spores of WT, *cmr*Δ3-ON, or Δ*cmrR*Δ*cmrT*. On day 2 p.i., cecal contents were collected for analysis using the NanoString nCounter Mouse Host Response Panel. **(A-C)** Volcano plots of differences in transcript abundance in mice inoculated with **(A)** WT, **(B)** *cmr*Δ3- ON, or **(C)** Δ*cmrR*Δ*cmrT* compared to mock-inoculated mice. Shown are Log_2_ fold changes in expression and adjusted p-values. Red and blue dots highlight genes with adjusted p-value ≤ 0.05 (above the horizontal dotted line). Red = genes with Log_2_ fold change ≥ 1; blue = genes with Log_2_ fold change ≤ −-1 (vertical dotted lines). **(D-E)** Venn diagrams with the number of differentially expressed (DE) genes unique to or shared among groups. Genes with **(D)** increased or **(E)** decreased differential expression in WT, *cmr*Δ3-ON, and Δ*cmrR*Δ*cmrT* relative to mock. **(F)** Heat map with Log_2_ fold changes of the shared DE genes.

## DISCUSSION

In this study, we sought to evaluate the relationship between colony morphology and the virulence of *C. difficile*. Our original approach entailed analyzing naturally arising, wild-type rough and smooth colony variants in a mouse model of CDI. These experiments suggested no difference in the ability of the variants to colonize the intestinal tract or cause disease. Further characterization of these infections revealed that inversion of the *cmr* switch, and by inference phase variation of CmrRST, occurred during infection. Specifically, independent of the starting orientation of the *cmr* switch, *C. difficile* in feces rapidly shifted to a predominantly *cmr-*OFF state after inoculation. Tissue-associated *C. difficile* exhibited a transient *cmr*-ON bias 1 day p.i., but showed similar proportions of *cmr*-ON/OFF as the fecal populations by day 2 p.i. These results undermined our comparison of rough and smooth colony variants in the infection model but revealed dynamic changes in the phenotypic makeup of *C. difficile* populations during infection.

As an alternative strategy, we evaluated phenotypically locked mutants that produce only rough (*cmr*Δ3-ON) or only smooth (Δ*cmrR*Δ*cmrT*) colonies *in vitro* in a mouse model of CDI (31). The *cmr*Δ3-ON or Δ*cmrR*Δ*cmrT* mutants showed no differences in weight loss, fecal burden, or toxin titers, though the *cmr*Δ3-ON-inoculated mice displayed greater colonic shrinkage than the mock-inoculated mice (37,38). In contrast, direct competition of these mutants in mice revealed that the Δ*cmrR*Δ*cmrT* mutant markedly outcompeted *cmr*Δ3-ON^R^, suggesting greater fitness of *cmr*-OFF *C. difficile*. These results are consistent with the phase variation dynamics of wild-type *C. difficile*, showing a shift to a predominantly *cmr-*OFF population during infection. A switch to a *cmr*-OFF bias was also observed in a hamster model of acute CDI and for Clade 5 *C. difficile* in a mouse model of CDI (26,43). Together these findings indicate that the *cmr*-OFF variant is more fit than the *cmr*-ON variant and suggest that *C. difficile* cells with smooth colony-associated phenotypes are better equipped to survive in the intestinal tract.

We also explored the host immune response through transcriptional changes in mice inoculated with WT, *cmr*Δ3-ON, and Δ*cmrR*Δ*cmrT*. NanoString analysis revealed 31 differentially expressed (DE) genes shared among all infected groups, and *cmr*Δ3- ON inoculated mice showed the highest Log_2_ fold-changes for most of these genes in comparison to WT and Δ*cmrR*Δ*cmrT*. Multiple genes identified, including *Lrg1* and *Lcn2*, were previously found to be upregulated during and associated with intestinal inflammation and infection (44,45). Expression of *Il33* was also higher in the *cmr*Δ3-ON inoculated mice; this alarmin has been shown to influence the severity of murine CDI symptoms (46). In addition, among the DE genes unique to *cmr*Δ3-ON inoculated mice, several are involved in the IL-17 signaling pathway, which has been determined to contribute to the severity of *C. difficile* infection in murine models of CDI (42,47–49). These differences in innate immune gene expression point toward a more robust immune response against the phenotypically locked rough mutant, and likely against *cmr*-ON cells more broadly.

The convergence towards *cmr-*OFF in WT *C. difficile* cells, the higher fitness of the Δ*cmrR*Δ*cmrT* mutant, and the stronger transcriptional response of mice to the *cmr*Δ3-ON mutant suggest preferential immune clearance of *cmr*Δ3-ON *C. difficile*. This effect may be driven by one or more of the phenotypes that distinguish the rough (*cmr*- ON) and smooth (*cmr*-OFF) colony variants (26,31,50). Recent work identified several CmrRST-regulated genes, including *mprA* and *mrpB*, involved in the production of elongated bacilli and cell chains observed in the rough variants (50). MrpA was found to interact with the cell division protein MinD leading to a model in which MrpAB production in *cmr*-ON cells disrupts proper cell division and the observed rough colony phenotypes. Increased cell size may enhance complement- and antibody-mediated pathogen neutralization, imposing a fitness cost to the *cmr*-ON state (51,52). Alternatively, impairment of cell division itself may compromise *cmr*-ON cell fitness during infection.

The advantage of CmrRST phase variation, and of the *cmr*-ON state specifically, remains unclear. The *cmrRST* locus, including the regulatory region upstream of the operon, is highly conserved in *C. difficile*, suggesting functional importance (26). The transient increase in the proportion of *cmr*-ON cells associated with tissue 1 day p.i. may indicate a niche in which this variant is advantageous. Prior work indicated distinct metabolic needs of the rough and smooth colony morphotypes, raising the possibility that nutrient availability impacts the relative fitness of the *cmr*-ON and *cmr*-OFF variants (13). A recent study showed that co-culture of *C. difficile* with *Enterococcus faecalis*, which often co-infects CDI patients, promotes the *cmr*-ON state *in vitro* (53–55). This finding suggests that microbe-driven environmental factors may underlie phase variation between *cmr*-ON and -OFF states. It is also possible that the *cmr*-ON state is advantageous outside of CDI, such as during asymptomatic carriage. Additional research is needed to elucidate the specific CmrRST-dependent factors and phenotypes, as well as host and microbial factors, that influence phase variation and shape the phenotypic makeup of *C. difficile* during infection and the severity of CDI.

## MATERIALS AND METHODS

### Bacterial growth conditions

The strains used in this study are listed in Table S1. Overnight cultures of *C. difficile* R20291 wild-type and mutant strains were prepared in either 3 mL of BHIS medium consisting of Brain Heart Infusion (Becton, Dickinson, and Company) supplemented with 5% yeast extract (Becton, Dickinson, and Company) or in TY medium consisting of Tryptone broth (Gibco Bacto) supplemented with 2% yeast extract and 0.1% sodium thioglycolate (Sigma). Cultures grew statically at 37°C in an anaerobic chamber (Coy Lab Products) with an atmosphere of 10% H2, 5% CO2, and 85% N2. *E. coli* strains were grown aerobically in Luria-Bertani (LB) broth or LB-agar (Fisher Scientific) statically at 37°C. Antibiotics were included in the media as appropriate at the following concentrations: 20 µg/mL chloramphenicol, 10 µg/mL thiamphenicol, 50 µg/mL kanamycin, 1200 µg/mL spectinomycin, and 250 µg/mL D-cycloserine.

### Construction of bacterial strains and plasmids

To create marked strains, an *aad9* spectinomycin resistance cassette from *Enterococcus faecalis* was integrated on the *C. difficile* chromosome between CDR20291_2492 and CDR20291_2493 via allelic exchange using the pMSR0 or pJB94 plasmids as previously described (56–59). Upstream and downstream homology regions were amplified from R20291 genomic DNA with primers R2914 + R3193 and R3192 + R2917, respectively, and cloned into BamHI-digested pMSR0 using Gibson assembly. A SphI site was added between the homology arms to allow insertion of DNA intended for integration on the chromosome, yielding pRT2891. The *aad9* cassette was amplified using primers R3416 + R3417 and cloned into SphI-digested pRT2891, yielding pRT3098. Clones were confirmed using colony PCR and Sanger sequencing with primers R2987 + R2988. To mark *cmr*Δ3-ON, the *aad9* cassette and flanking homology arms for integration between CDR20291_2942 and CDR20291_2943 were amplified from pRT3098 by PCR using R3951 + R3952. The PCR amplicon was cloned into the SpeI and XhoI sites of pJB94 by digestion and ligation, generating pRT3371. The final plasmids, pRT3098 and pRT3371, were introduced into heat-shocked *C. difficile* R20291 and *cmr*Δ3-ON, respectively, via conjugation with *E. coli* HB101(pRK24) as described previously (29,60,61). Mutants were confirmed through whole-genome sequencing.

### Colony morphology microscopy

*C. difficile* strains were streaked onto BHIS agar supplemented with 0.1% L-cysteine and incubated at 37°C for 48 hours before imaging. Plates were removed from the anaerobic chamber, and colonies were imaged at 2X magnification using the Keyence BZ-X810 microscope.

### Spore purification

*C. difficile* growth containing spores was collected from 70:30 agar after 72 hours of growth, suspended in 10mL DPBS (Gibco ^TM^), and stored aerobically for sucrose gradient purification (62,63). Spores were washed five times in DPBS + 1% BSA and stored at room temperature until inoculation.

### Animal experiments

Mouse experiments were performed under the guidance of veterinary staff within the UNC Division of Comparative Medicine (DCM). All animal studies were done with prior approval from the UNC Institutional Animal Care and Use Committee and in compliance with guidelines from the American Veterinary Medical Association.

Groups of male and female C57BL/6 mice (Charles River Laboratories, age 8- to 10-weeks) were given a previously described antibiotic regimen to render the mice susceptible to *C. difficile* (35). A cocktail of kanamycin (400 mg/L), gentamycin (35 mg/L), colistin (850,000 units/L), vancomycin (45 mg/L), and metronidazole (215 mg/L) was provided in their water *ad libitum* 7 days before inoculation for 3 days, then the mice were returned to regular water. A single intraperitoneal dose of clindamycin (10 mg/kg body weight) was administered 2 days prior to inoculation. Mice were randomly assigned to groups and then were inoculated with a total of 10^5^ spores by oral gavage. Mock-inoculated mice were included as controls, and cage changes were performed every 48 hours post-inoculation (p.i.). Animal weights were recorded, and fecal samples were collected every 24 hours p.i. Fecal samples were homogenized, and dilutions (1:10 in DPBS) were plated on TCCFA plates, which contain 0.1% of the germinant taurocholate, cefoxitin (10 mg/mL), D-cycloserine (25 mg/mL), fructose (6 g/L), and agar (20 g/L). On specified days p.i., subsets of mice from each group were euthanized by CO_2_ asphyxiation followed by cervical dislocation. After euthanasia, the large intestines were collected to measure colon length (cm) and enumerate CFU, cecal contents were collected for measurement of toxin levels, and cecal tissue was stored in RNA*later* Solution (Invitrogen) at −20°C.

For CFU quantification in tissues, the contents of the large intestines were carefully removed, and the tissues were rinsed with DPBS. After sectioning into cecum, proximal, and distal colon segments, the tissues were treated with 0.1% dithiothreitol (DTT) in DPBS, rocking at room temperature for 30 minutes to dissociate the mucus layer. The remaining tissue was placed in DPBS and homogenized (bead beater, 3 minutes), then serial dilutions were plated on TCCFA.

For competition experiments, spores from wild-type (WT^R^) or *cmr*Δ3-ON^R^ spectinomycin-resistant mutants were combined at a 1:1 ratio with wild-type (WT) or Δ*cmrR*Δ*cmrT* spectinomycin-sensitive strains, respectively. Antibiotic-treated mice were inoculated with 10^5^ spores total. Animal weights were recorded, and fecal samples collected every 24 hours p.i. were homogenized and plated on TCCFA with and without 1200 µg/mL of spectinomycin (Sigma). To calculate the competitive index (CI), the ratio of resistant to sensitive bacteria for each fecal sample (resistant CFU/[total CFU – resistant CFU]_output_) was divided by the ratio of resistant to sensitive bacteria in the initial spore inoculum (resistant CFU/[total CFU - resistant CFU]_input_).

### Competition experiments *in vitro*

The *cmr*Δ3-ON^R^ and Δ*cmrR*Δ*cmrT* mutants were grown in TY broth overnight. The cultures were normalized to an absorbance (OD_600_) of 1.0 and mixed at a 1:1 ratio. The co-cultures were diluted (1:50) into fresh TY daily, and CFU were enumerated on BHIS-agar with and without 1200 µg/mL of spectinomycin. The competitive index was calculated as described above.

### Quantification of switch orientation by quantitative PCR

To lyse *C. difficile*, spores, tissue, and fecal samples were treated with lysozyme and subjected to bead beating The genomic DNA was purified by phenol:chloroform:isopropanol (25:24:1) extraction and precipitated in ethanol (23,60). Quantitative PCR (qPCR) was performed as previously described, with 100 ng of DNA per 20 µL-reaction, 100 nM of each primer, and SensiMix SYBR and Fluorescein Kit reagent (Bioline) (25,31). Reactions were run on LightCycler® 96 Instrument (Roche) with the following three-step cycling conditions: 95°C for 10 minutes, followed by 40 cycles of 95°C for 30 seconds, 60°C for 1 minute, and 72°C for 30 seconds. Data were analyzed as described previously using *rpoA* as the reference gene (25). Primers used are listed in Table S2.

### Vero cell rounding assays

We used previously described protocols to quantify toxin production by *C. difficile* (23,64). Briefly, 5 ×10^4^ Vero cells in 90 µL were seeded in tissue-culture treated 96-well plates and incubated overnight at 37°C and 5% CO2. The following day, mouse cecal contents were thawed at room temperature, weighed, and suspended in DPBS to make an initial 1:10 dilution. The supernatants were filtered-sterilized (0.45 µm) and serially diluted (1:10) in DPBS on ice. Dilutions (10 µL) were added to the Vero cells and incubated overnight (∼ 18 hours) at 37°C. Cell rounding was assessed using light microscopy at 10X magnification. Toxin titers were calculated as the reciprocal of the highest dilution causing over 80% cell rounding, normalized to the grams of cecal contents. Samples from mock-inoculated animals were included as negative controls.

### Mouse quantitative reverse transcription PCR

Tissue stored in RNA*later* was thawed at room temperature, and total RNA was purified using the RNeasy Mini Kit (Qiagen) followed by treatment with RNase-Free DNase (Qiagen) per manufacturer’s instructions. The cDNA was synthesized using M-MuLV Reverse Transcriptase (NEB), Random Primer Mix (NEB), and RNase Inhibitor (NEB) as instructed by the manufacturer. Transcript abundance was measured by qRT-PCR using 100 ng cDNA, primers at a final concentration of 500 nM, and SensiMix SYBR and Fluorescein Kit (Bioline). Reaction parameters were set as described above, except with an annealing temperature of 55°C. Primers used for *Il1β*, *Cxcl1*, *Lcn2*, and *Tnfα* are listed in Table S2. The housekeeping gene *Tbp* was used as the reference gene, and the data are normalized to the mean value for the mock controls.

### NanoString analysis

RNA was extracted from cecal tissue as described above. Samples were submitted to the UNC Biospecimen Processing Facility for quality assessment using the 4200 TapeStation. RNA was then submitted to the Duke Microbiome Center for transcriptional analysis using the nCounter Mouse Host Response Panel. Raw data were imported into nSolver Advance Analysis software (v 4.0) for normalization and differential expression analysis. Correction for multiple comparisons was performed using the method of Benjamini-Hochberg (65).

### Statistical analysis

With the exception of NanoString analysis, all statistical tests were performed in GraphPad Prism (10.3.1). *C. difficile* CFU, mouse weights, toxin titers, colon length, and qRT-PCR data were analyzed using one-way ANOVA with Kruskal-Wallis test and Dunn’s multiple comparisons. A p-value of < 0.05 was considered statistically significant.

## Acknowledgements

We thank Duke University School of Medicine for the use of the Microbiome Core Facility, which provided Gene Expression service. This work was supported by grant R01AI143638 to R.T. from the National Institute of Allergy and Infectious Diseases. The funders had no role in study design, data collection and analysis, decision to publish, or preparation of the manuscript.

